# Spatio-temporal structure and reproductive success in a rook (*Corvus frugilegus*) colony

**DOI:** 10.1101/409664

**Authors:** Orsolya Feró, Miklós Bán, Zoltán Barta

**Affiliations:** MTA-DE “Lendület” Genome architecture and recombination laboratory, University of Debrecen, H-4010 Debrecen, Egyetem tér 1, Hungary.; MTA-DE Behavioural Ecology Research Group, Department of Evolutionary Zoology and Human Biology, University of Debrecen University of Debrecen, H-4010 Debrecen, Egyetem tér 1, Hungary.

**Keywords:** coloniality, colony structure, colony formation, breeding success, clutch size, hatching success, nest density, nest site selection, colony syncrony

## Abstract

We have studied how individual decisions about timing of breeding and nest location affected the reproductive success of rooks by tracing the formation of a rook breeding colony in Hortobágy NP (Hungary), during the breeding season in 1999.

We have found that birds who built nests earlier also laid eggs earlier and had larger clutch size but had no had more offspring than late nesters. Distance of the nest from the centre or edge of the colony did not affect the reproductive success though rooks generally settled closer to the centre and further from the edge than it can be expected by assuming random distribution. The colony showed high breeding synchrony since date of egg laying varied less than date of nesting but synchrony did not influence the breeding success. Hatchling’s survival rate and number of offspring increased with local nest density and rooks clustered their nests more than it can be expected by random settlement. Nest sites of early and late nesters did not differ regarding the distance from the centre or the edge of the colony but late nesters chose nest sites in more densely populated regions.

The results indicate that individuals may follow different strategies to increase their reproductive success. It is prospectively advantageous to build the nest and lay eggs earlier, in turn, it’s worth nesting in already more densely nested areas for late beginners. We suggest that a game theoretical approach may be useful if we try to understand the adaptive significance of colonial breeding.

## INTRODUCTION

The breeding assembles of birds has attracted considerable attention from ornithologists and behavioural ecologists over the last decades (e.g. Lack 1968, Wittenberger & Hunt 1985, Danchin & Wagner 1997). During this long course of investigation researchers have identified many possible costs and benefits associated with colonial breeding. For instance, colonial breeding may increase ectoparasite infection (Brown & Brown 1986), increase competition for food, nesting sites, nesting material and mates, and even lead to kleptogamy (for a review, see Wittenberger & Hunt, 1985). The advantages may include decreased predation risk (Hoogland & Sherman 1976, Wiklund & Andersson 1994, Perry & Andersen 2003), increased foraging efficiency (information center hypothesis - Ward & Zahavi 1973, Brown 1988, Barta & Szép 1992, 1995, Barta & Giraldeau 2001; local enhancement hypothesis - Buckley 1997, Pöysa 1992; recruitment center hypothesis - Evans 1982, Richner & Heeb 1996) or increased extra-pair mating (Hoi & HoiLeitner 1997). It is still not clear which of these factors is mainly responsible for the evolution and maintenance of colonial breeding. It is probable, as Danchin and Wagner (1997) indicate, that colonies emerge as the result of multiple, complexly interacting factors, which vary according to species or populations.

The evolutionary explanation of colonial breeding is further complicated by the fact that these costs and benefits may change not just between species or populations, but between individuals, either because they occupy different spatial (e.g. densely vs. rarely populated) and temporal (early vs. late breeders) position within the colony or are of different types (e.g. dominants vs. subordinates). Nesting in more densely populated locations can be beneficial e.g. by increasing the possibility of information exchange about the location of food (Brown 1988) or decreasing the rate of predation (Hatchwell 1991, Hotker 2000) but, on the contrary, high density sometimes increases the risk of predation (Clode 1993, Bellinato & Bogliani 1995). Further costs can be increased rate of parasite infection (Tella 2002), intraspecific aggression (Stokes & Boersma 2000, Hotker 2000) or competition (Hoogland & Sherman 1976). Reproductive success of birds occupying central nest sites can also differ from those that nest in the periphery (Coulson 1968, Brunton 1997). Early breeders can occupy better nest sites and achieve higher breeding success (Burger & Shisler 1980). They and their offsprings can also have more time to improve their physical condition and prepare for the subsequent winter, so their survival rate can be higher as well (Hussell 1972). However, reproductive success can be influenced more by the timing of breeding relative to the others (syncrony) than the exact date (Hatchwell 1991, Murphy & Schauer 1996). Moreover, different individuals (according to age, experience, condition) may follow different strategies to achieve higher reproductive success (Sasvári & Hegyi 1994).

It is likely that these factors lead to that distinct parts of the colonies provide various benefits and costs for different individuals. It is expected that under natural selection individuals act to maximize their success by occupying better nest sites within the colony and by breeding at optimal time. This task, however, is far from trivial because the best place and time depend not just on the environmental characteristics of a site, but also on the behaviour of other individuals in the population. For instance, let us assume that it is best to breed early in the most densely populated part of the colony. But how can an early breeder decide which will the most crowded part of the breeding assembly when this is dependent on the decision of late nesters? Consequently, the formation of the spatio-temporal structure of colonies can only be understood in a game theoretical context which, in turn, might provide a better basis for investigating the adaptive significance of colonial breeding.

Until recently only a few studies investigated how a colony is formed as a results of individual decisions. For instance, Burger & Shisler (1980) found that in Herring Gull (*Larus argentatus*) colonies one or a few epicentres are occupied first and late breeders have no choice but nest in the less advantageous edge of the colony. Alternatively Velando & Freire (2001) showed that colony formation of the European Shag (*Phalacrocorax aristotelis*) fits the so-called central-satellite model. This model suggests that low-quality late breeders build their nests close to high-quality early-breeder birds, which do not necessarily nest in the centre of the colony. This type of distribution might provide higher chance to obtain a better breeding site or mate for the low-quality birds in the following season and, for the high-quality birds, higher chance of extra-pair mating.

Our study’s aims were to observe the formation of a Central-European rook breeding colony during the breeding season and explore whether individual reproductive success is affected by the timing of breeding (early and late breeders, breeding synchrony) or by the location of nest (central or peripheral) or local nest density (number of neighbours). We also investigated whether rooks have any preference in choosing nest sites and how these choices influence their reproductive success.

## METHODS

The fieldwork was carried out in Hortobágy-Szálkahalom (21°14′E, 47°34′N), district of Hortobágy National Park, Hungary, during the breeding season of 1999. A large stable rook breeding colony of 800-900 nests exists there occupying a plantation of Black Locust (*Robinia pseudoacacia*).

On the basis of the preliminary observations in the preceding year, we know that in this colony rooks begin to build new nests or renovate the old ones in February. To follow the formation of the colony we counted the nests at 11 sampling dates (every 5-12th day) from February to April and individually marked all trees with nest(s). These individual marks were then used to identify the nests. If there were more than one nest on a tree we also used digital photos for nest identification. We recorded 1115 nests on 555 marked trees during the breeding season.

After every field-day we chose randomly 15% of the newly recorded nests for further studies (165 nests in total) – this was the amount, according to the preliminary observations, that we were able to check frequently, without disturbing the colony too much. A long aluminium pole with a perpendicularly fixed mirror on its end was used to observe the content of the chosen nests. After we found eggs in a nest we monitored that nest every 3-4th day. When the chicks began to hatch we calculated the laying date of the first egg (elapsed days from the 31th of January) by estimating the age of the first chick and by using the 17 day average (Roskaft 1983b) as incubation length. We recorded the clutch size (maximum number of laid eggs) and brood size for the selected nests. We chose the number of 7-10 day old offspring as an estimate for brood size because the most of the nestlings’ mortality occur in the first 10 days after the hatching (Roskaft et al. 1983). The quotient of the brood size and the clutch size was also calculated for the observed nests and we refer to this as offspring’s survival rate later in this paper.

The nests that were destroyed by vernal high wind or because of some other accident and those that were abandoned and disappeared before completion (25 nests altogether) were excluded from the analyses that dealt with the reproductive success. We could not always gather all information about the nests, sometimes the nest could not be positively identified or we were unable to reach it, and it was often difficult to estimate the number of offspring exactly. However, we used as much data as possible for each test, so (excluding the doubtful data) the sample sizes varied between analyses.

To obtain the position of nests within the colony we made an accurate map of the trees with nest(s) by the distance measurement method in autumn far after the breeding season in order to reduce disturbance (for the method, see Boose et al. 1998).

Using the relative co-ordinates of the trees we calculated several spatial variables to characterize nests’ position. We determined the nests’ distances from the centre of the colony and from the edge of the colony. We used the centre of gravity of all nests as centre and the smallest convex polygon that included all of the nests at the edge. Number of neighbouring nests within 6 metres at the date of hatching were also counted for all nests. Usually the distance of the nearest neighbour or average distance of several neighbours is used for such investigations but the number of close neighbours may be more appropriate and more important regarding the costs and advantages of group living (e.g. Barta & Giraldeau 2001). We chose the 6 metre range because the average distance between nests was around 4.5 metres and we wanted to describe the effect of the closest nest surroundings within which social interactions may have the strongest effect. To find out whether the results are dependant on the selected distance we also made the analyses using 9, 12 and 15 metre ranges.

Relative syncrony was also determined as the difference between each chosen nest’s hatching date and the median of hatching dates in days.

To find out what preferences rooks may have and whether they try to minimize the costs or rather exploit the benefits of group living when they choose their nest site, we selected 100 random points within the edge of the colony for each sampling date and compared them to the newly built nests at the given date in respect of the positions of the nests, i.e. distance from the centre, distance from the edge and local nest density (number of neighbouring nests). We also tested whether the preference changed during the season, so if there were any changes in the position parameters of the observed nests compared to the randomly selected points.

We used Spearman rank correlations and Mann-Whitney tests for statistical analyses with two-tailed significance level.

## RESULTS

### Starting time, colony synchrony and breeding success

Nests were built from the beginning of February to mid April while offspring hatched from the beginning of April to May. The start of nest building varied more in time than egg laying (Levene statistic: 18.987, df_1_ = 1, df_2_ = 231, P < 0.001; Fig. 1).

**Figure 1.**
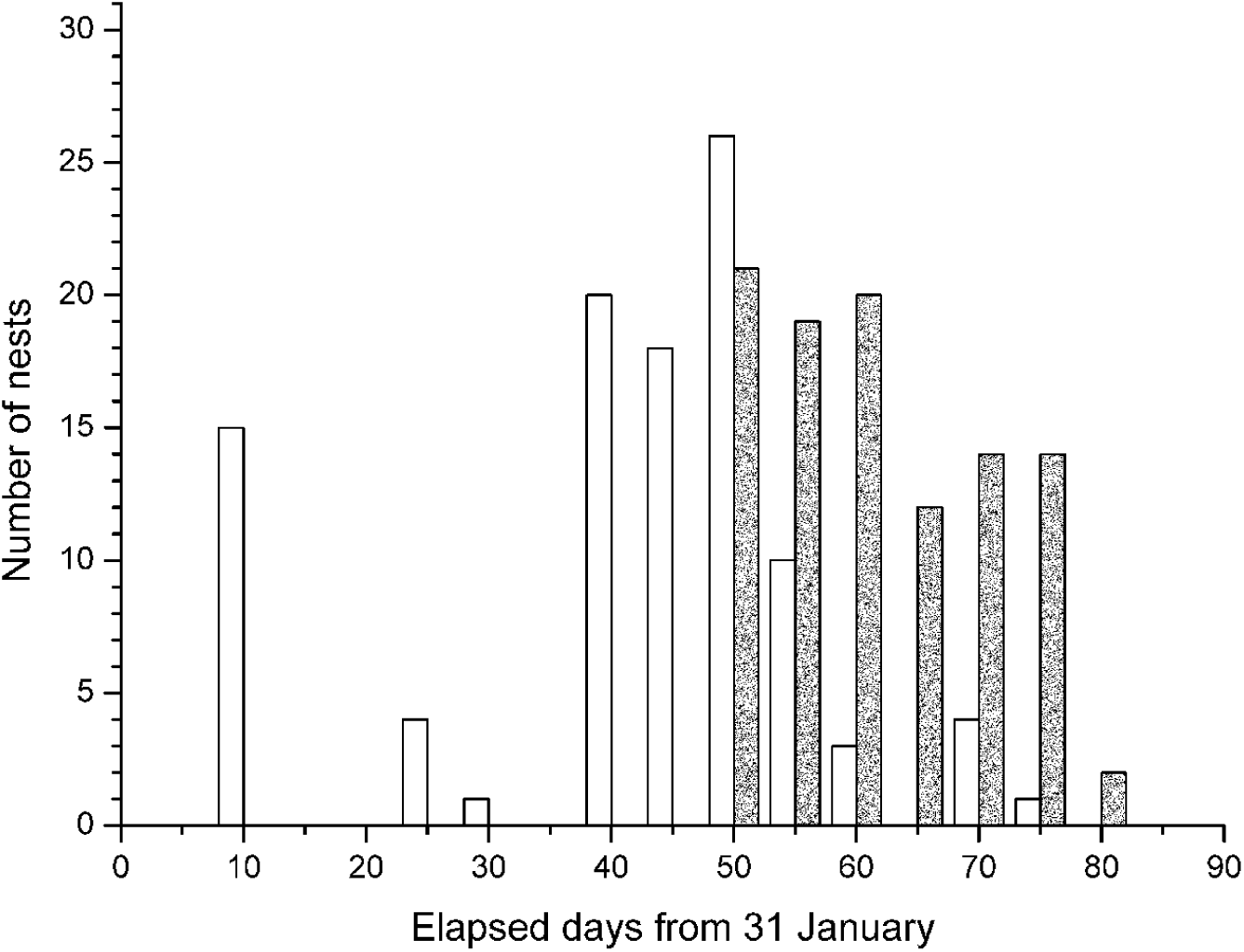
Temporal distribution of nest building and egg laying. Bars represent the number of nests that were started to build (white) and number of nests in which the first egg was laid (grey) grouped into five-day periods. n = 20, mean ± SD: 40.00 ± 5.94 and 76.33 ± 8.80 days respectively.

Rooks that had started to nest earlier also laid their eggs earlier (r_s_ = 0.610, n = 102, P < 0.001) but they waited much more between nesting and egg laying than those who started later (r_s_ = -0.695, n = 102, P < 0.001; Fig. 2).

**Figure 2.**
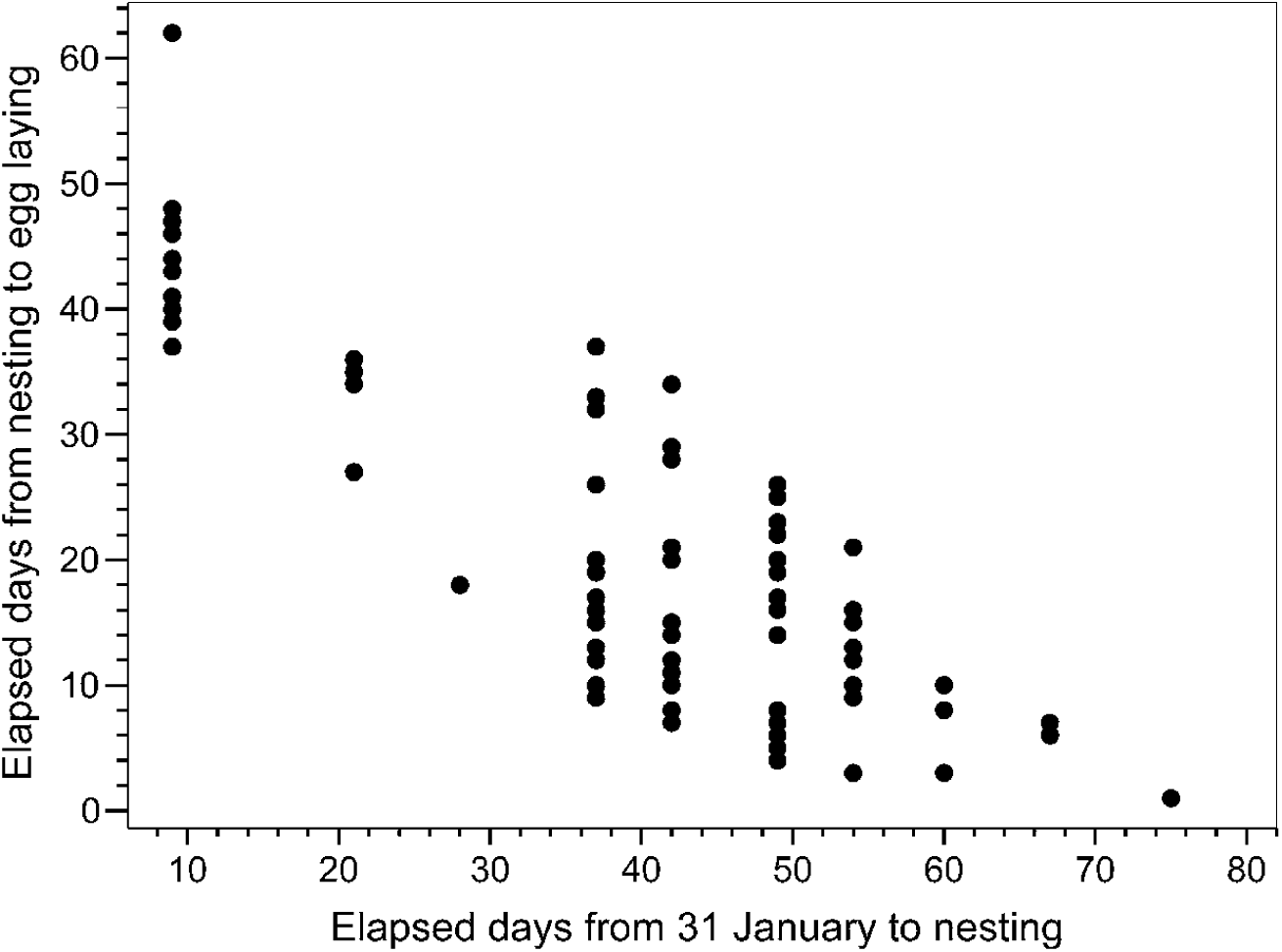
Relationship between the date of nesting and the elapsed time from nesting to egg laying. Spearman rank correlation: r_s_ = -0.695, n = 102, P < 0.01.

There was a negative correlation between the date of nesting (and date of egg laying) and clutch size, so rooks that started earlier laid more eggs, but the date of nesting or egg laying had no effect on their brood size (Table 1). There was no significant difference in hatchling’s survival rate either between early and late nests or between early and late clutches (Table 1). Relative synchrony influenced neither the brood size nor hatchling’s survival rate (Table 1).

**Table 1.**
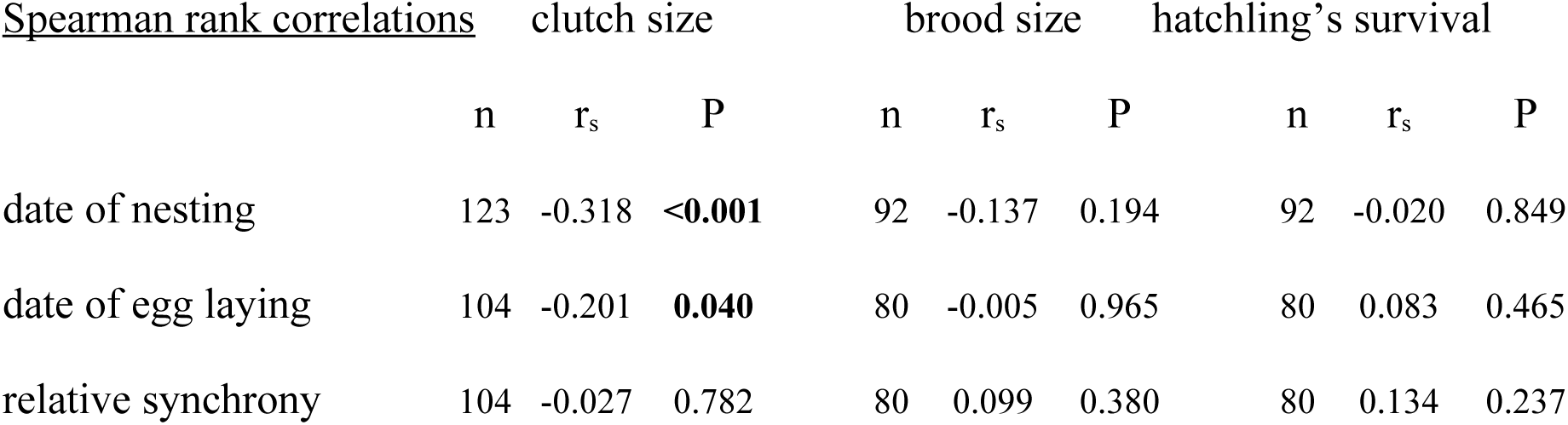
Statistical results for the correlations between timing of breeding and reproductive success. Dates are elapsed days from 31 January and relative syncrony is the difference between the nest’s hatching date and the median of hatching dates in days.

### Position and breeding success

Neither clutch size nor number of hatchlings correlated with distance from the centre or the distance from the edge of the colony (Table 2). We did not find any relationship between hatchling’s survival rate and either the distance of the nest from the centre or the edge of the colony (Table 2).

**Table 2.** Statistical results for the correlations between nest location, nest density and reproductive success. Distances are given in metres; number of neighbouring nests were counted for each nest at the date of hatching.

| <u>Spearman rank correlations</u> | clutch size |  |  | brood size |  |  | hatchling's survival |  |  |
| --- | --- | --- | --- | --- | --- | --- | --- | --- | --- |
|  | n | r<sub>s</sub> | P | n | r<sub>s</sub> | P | n | r<sub>s</sub> | P |
| distance from centre | 125 | -0.061 | 0.497 | 95 | 0.054 | 0.060 | 95 | 0.108 | 0.298 |
| distance from edge | 125 | 0.034 | 0.710 | 95 | 0.070 | 0.502 | 95 | 0.003 | 0.979 |
| number of neighbours |  |  |  |  |  |  |  |  |  |
| < 6 metres | 103 | -0.136 | 0.170 | 80 | 0.262 | **0.019** | 80 | 0.382 | **<0.001** |
| < 9 metres | 103 | -0.164 | 0.098 | 80 | 0.208 | 0.064 | 80 | 0.311 | **0.005** |
| < 12 metres | 103 | -0.148 | 0.135 | 80 | 0.233 | **0.037** | 80 | 0.347 | **0.002** |
| < 15 metres | 103 | -0.124 | 0.213 | 80 | 0.225 | **0.045** | 80 | 0.332 | **0.003** |
| on the same tree | 126 | 0.171 | 0.056 | 95 | 0.105 | 0.310 | 95 | 0.066 | 0.527 |

Hatchling’s survival rate positively correlated to the number of neighbouring nests within 6 metres at the date of hatching so where the local nest density was higher the survival rate was also higher and more hatchling were hatched in these nests as well (Fig. 3, Table 2). There was no correlation between clutch size and number of neighbours (Table 2). The results were similar if we changed the range; hatchling’s survival rate and brood size (except for the 9 metre range) increased with number of neighbouring nests, but clutch size did not depend on the number of neighbours (Table 2). It was not improbable that breeding success primarily depended on the characteristics of the tree and the trees offering better nest sites attracted more occupants. Therefore we analysed the correlation between the number of nests on the same tree at the date of hatching and breeding success, but we did not find any relationship between them (Table 2). The number of nests on the occupied trees varied between 1 and 6 nests per tree and was 1.6 in average.

**Figure 3.**
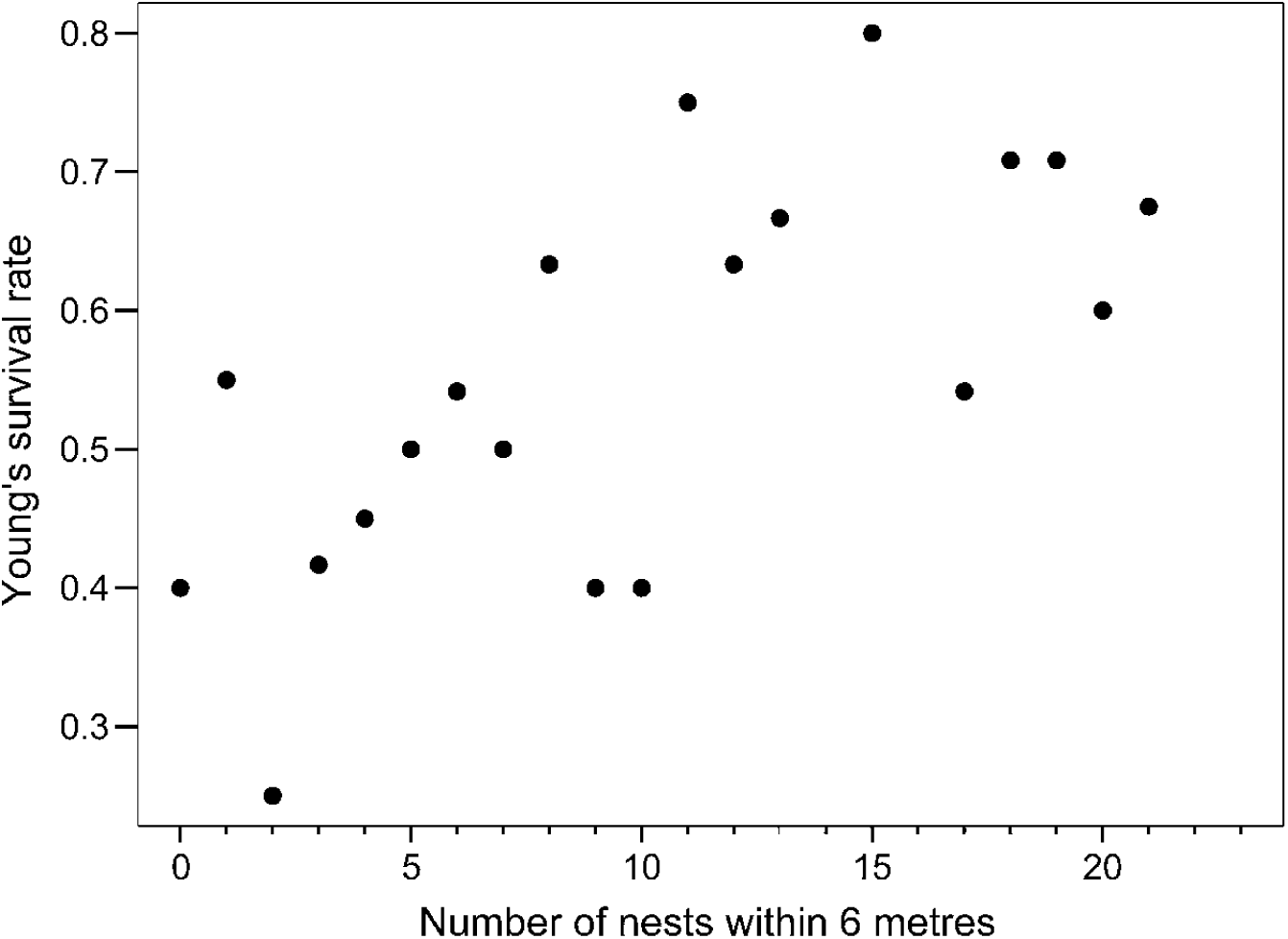
Relationship between local nest density (number of neighbouring nests within 6 metres) at the date of hatching and hatchling’s survival rate (medians). Spearman rank correlation: r_s_ = 0.382, n = 80, P < 0.001.

### Spatio-temporal pattern

Each sampling date newly build nests appeared in more densely nested areas than the randomly selected points (Fig. 4). The nests were built closer to the centre and farther from the edge of the colony compared to the random points though the difference was not always significant during the season (Fig. 4).

**Figure 4.**
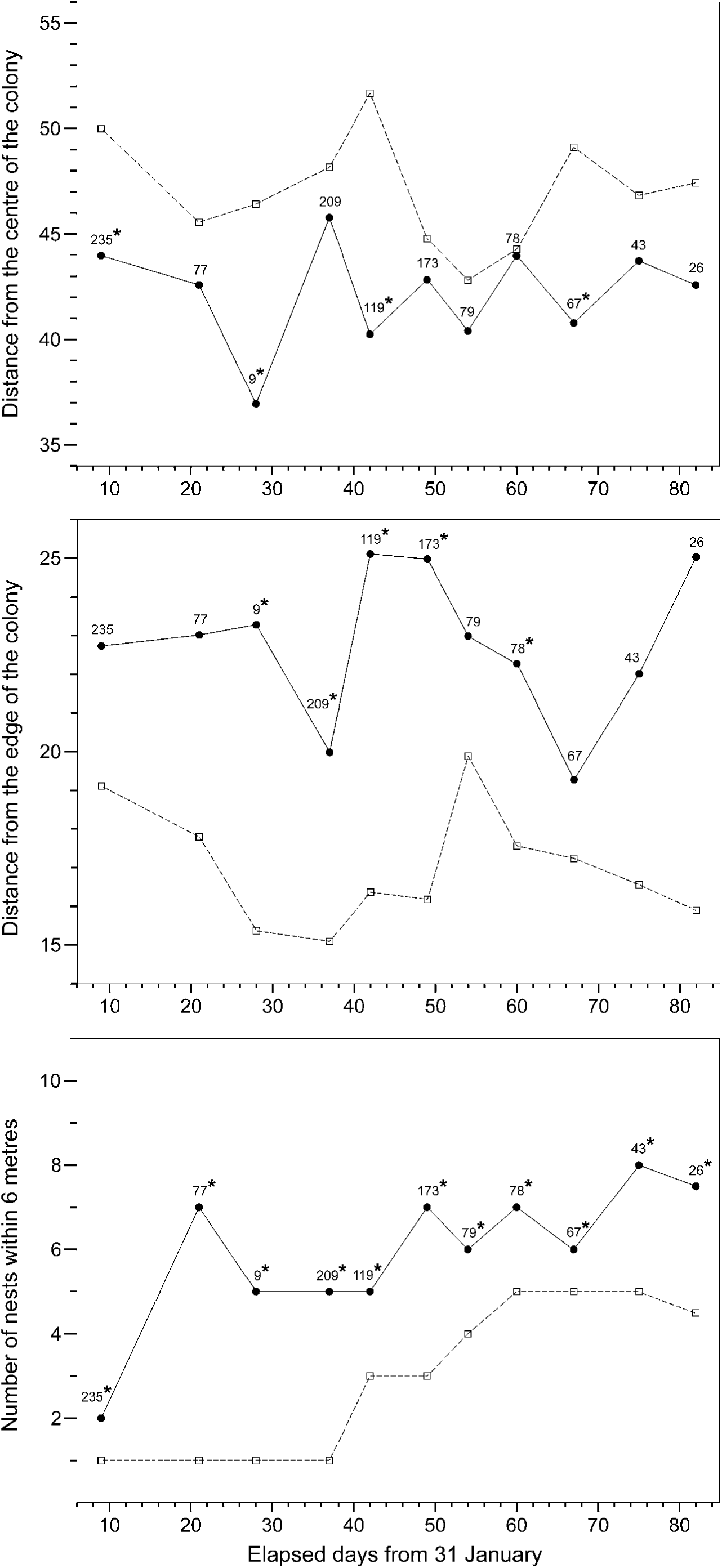
Distance of the newly built nests from the centre of the colony, from the edge of the colony (metres) and number of neighbouring nests within 6 metres (medians, full circles) and those of the randomly selected points (medians, empty squares) at each sampling date. 100 random points were selected for each date, sample sizes of the observed nests are shown above the medians. In the marked (*) cases the values of the observed nests differed significantly from the values of the random points (Mann-Whitney test, P < 0.05).

Preference of early and late nesters did not differ in choosing periferal or central nest position as date of nesting correlated neither to distance from the centre (r_s_ = -0.014, n = 1115, P = 0.647) nor distance from the edge (r_s_ = 0.009, n = 1115, P = 0.756). Late nesters, not surprisingly, had more close neighbours at the date of nesting than early nesters (r_s_ = 0.410, n= 1115, P < 0.001).

## DISCUSSION

In the observed rook colony in Szálkahalom we found that reproductive success of individuals are different in relation to timing of breeding and nest position.

Breeding success was mainly affected by the number of close neighbouring nests; more hatchling had survived the first week where the local nest density was higher. Although nesting in the most densely populated areas are often disadvantageous in some colonially breeding species e.g. because of increased intraspecific aggression (Stokes & Boersma 2000, Hotker 2000) or competition (Hoogland & Sherman 1976), it may be worth living in neighbourship to rooks. Probably it is because rook breeding colonies serve as information centres about the location of food. Rooks usually feed in loose flocks around the colony (e.g. Patterson et al. 1971). During the hatching period and for 2-4 weeks after hatching, female rooks stay at the nests and they and the offspring are fed by the males (Roskaft 1983b). The female’s condition, and hence the hatching rate and the survival of offspring is mainly affected by the male’s foraging efficiency during this period (Roskaft 1983b, Roskaft et al. 1983). Rate of information exchange (by monitoring the neighbours’ foraging success) thus individual foraging efficiency (by following successful foragers to the food patch) is presumably higher where local nest density is higher therefore more peers are present (Ward & Zahavi 1973, Barta & Giraldeau 2001).

Another possibility is that rate of extra-pair copulation is higher in more densely populated areas. This may reduce the number of infertile eggs so increase hatching rate as Brown & Brown (2001) suggested and, if high quality males have higher chance for extra-pair copulation (EPC), the offspring can be of better quality and have higher survival rate. Promiscuity is not rare among rooks, but most EPC attempts involve incubating, thus already fertilized females (Roskaft 1983a).

It is not likely that higher reproductive success in more densely nested areas resulted from lower predation risk. Decreasing effect of higher density on predation risk is not trivial, some studies support it (Hatchwell 1991, Hotker 2000) and others do not (Clode 1993, Bellinato & Bogliani 1995). Furthermore it is not the predation rate that determine hatchling’s survival in the rook (Roskaft et al. 1983). Breeding success of central and peripheral nesters are different in some species, which is usually explained by different predation risk (e.g. Brunton 1997), but we found that breeding success did not depend on the position of the nest.

Early breeders had significantly larger clutch size but this was not reflected later in their brood size and there was no difference in hatchling’s survival rate between early and late breeders. Since rooks (in areas where they are resident) have a strong association with their breeding colony throughout the year (Patterson et al. 1971), late nesters may not have considerable disadvantage because of fewer simultaneously breeding bird. A possible explanation of the relation between date of egg laying and clutch size is that older birds lay eggs earlier (Roskaft et al. 1983), assuming that young unexperienced birds do not lay clutches as large as older pairs do. Relative synchrony did not influence the reproductive success either, but the fact that date of breeding varied far less in time than date of nesting might indicate that breeding synchrony is important for rooks.

We found that rooks usually built their nests farther from the edge or closer to the centre of the colony and always aggregated their nests more than it would be expected in the case of random distribution. In their study, Brown & Brown (2000) investigated whether Cliff Swallows maximized nearest-neighbour distances of the nests during the settlement into colonies by comparing the observed nearest-neighbour distances to the expected distances assuming maximal spacing among nests. They found that birds generally settle closer to each other than the calculated maximized distances so they claimed that it is advantageous for Cliff Swallows to nest close to conspecifics. It might be more appropriate to determine how the observed distribution differs from the random if we want to investigate the effect of social interactions on nest site selection. If it is advantageous to nest close to the others, birds may build their nests closer to each other than in the case of random settlement. If this is costly, then they can maintain grater distance between nests than they would do if they settled randomly, though we may find that the observed nest-distances significantly smaller than the calculated maximized distances.

Since breeding success in the studied colony mainly depended on the local nest density at the date of hatching, late nesters might be provided an opportunity to estimate nest density accurately and increase their success by choosing already densely nested locations. Roskaft (1985) found that rooks in poorer condition laid eggs later and had lower hatching success. Maybe these rooks can compensate their disadvantage by exploiting the benefits of nesting in more densely populated areas. The following questions then arise: 1) why do early breeders begin to nest two months before they lay eggs? and 2) why do they not space their nests closer to each other? Early nesters may choose nest sites where more stable nests can be built (that can resist vernal high winds), higher nest sites that are more protected from occurrent persecution or nests that have survived from the year before. In addition rooks in better condition can afford to start building their nest early and defend it longer. They may gain relatively less benefit from social interactions (e.g. increased foraging efficiency) than by occupying the nests sites of better quality. It is worthy of note that early nesters also bred earlier so they and their offspring may have more time to prepare for the subsequent winter hence their survival rate may be higher. Early breeders have also more chance for a second brood if the first brood fails for whatever reason (Hussell 1972).

In summary, breeding colonies may have various advantages and costs for their members in relation to the individuals’ attributes (e.g. age, sex, experience, quality), the current environmental factors (available nest sites, weather, food abundance and distribution, etc.) and to the behaviour of all the colony members. Birds’ decisions about when and where to breed are based on various factors and there are alternate tactics that birds can follow to increase their breeding success. In our study, for example, reproductive success of an individual is prospectively higher if it begins to nest and lay eggs earlier, probably because early nesters may occupy better nest sites, furthermore, their and their offspring’ survival rate for the next year may be higher by breeding earlier. Later nesters can estimate the prospective nest density more accurately and increase their nestlings’ survival by nesting in a densely populated area and gain more benefit from social interactions.

## ACKNOWLEDGEMENTS

We are grateful to Zsolt Végvári for giving additional information about the studied colony and for his field assistance. We also thank András Kosztolányi for his severe comments on the first version of the manuscript and András Liker for further valuable notices. The fieldwork was allowed by a permission of the Hortobágy National Park. Our study was supported by an OTKA (Hungarian Scientific Research Fund) grant (T030343) to Z.B.

